# Human Mediodorsal Thalamus in Seizure Propagation

**DOI:** 10.64898/2025.12.01.691640

**Authors:** Olivia Marais, Chipi Lozano, Aida Risman, Masaya Togo, Sofia Pantis, Eric van Staalduinen, Lindsay Liu Yang, Robert Fisher, Vivek Buch, Josef Parvizi

## Abstract

**Background:** How different thalamic sites are recruited during seizure propagation remains poorly understood. Simultaneous recordings from multiple thalamic sites in patients with focal seizures provide a rare opportunity to investigate the spatiotemporal pattern of thalamic involvement during human epilepsy.

**Objective:** To characterize the recruitment patterns of mediodorsal (MD) thalamic subregion during seizures and their generalization to the contralateral hemisphere.

**Methods:** We analyzed 119 seizures from 23 patients (12 male, age range: 20-57y) undergoing multisite thalamic recordings. In accordance with current clinical standards, we determined the spatial and temporal features of thalamic seizure activity by visually reviewing intracranial EEG recordings from different seizure types in each individual patient.

**Results:** The procedure of multisite thalamic recordings had no complications. In total, we captured seizures originating from temporal lobes (63%), orbitofrontal (11%), frontotemporal (8%), occipital (8%), lateral frontal (4%), parietal (3%), and cingulate (2%) regions. Seizures were focal (76% in 21 patients), focal-to-bilateral tonic–clonic (FBTC, 9% in nine patients), or only electrographic (15% in six patients). Thalamic engagement was seen in 100% of patients occurring typically early during seizure evolution (83% within 15 seconds of seizure onset). Majority of FBTC seizures (73%) had faster thalamic recruitment, often within the first 5 seconds. The pulvinar (PLV) subregion was the most common first-activated thalamic site, particularly in temporal lobe seizures. Although the MD was involved in most seizures (88.2%), it was rarely the initial or sole thalamic structure engaged and more often followed anterior (ANT) and/or PLV sites. Contralateral propagation occurred in 66% of seizures and was strongly linked to MD involvement: the ipsilateral and contralateral MDs were engaged in about 95% of these cases. When ipsilateral MD engagement was absent, contralateral spread of seizures was uncommon. In majority of seizures (60%) that generalized to the contralateral hemisphere, the ipsilateral MD was involved before or simultaneously with the contralateral cortical sites. Importantly, seizures that first activated the MD originated mainly from the medial temporal lobes, whereas those spreading primarily to the contralateral cortex were mostly neocortical in onset.

**Conclusions:** The thalamic MD subregion was often involved after the other thalamic sites, but the MD sites, along with the massa intermedia connecting the two thalami, were significantly involved when seizures spread to contralateral hemisphere. Our findings suggest that a single thalamic lead capturing both MD subregions may yield important clinical information about laterality, origin, and generalization of seizures.

## Introduction

The thalamus has emerged as an important target for neuromodulation therapies in epilepsy.^1-3^ Deep brain stimulation (DBS) of the anterior nuclei of the thalamus is proven to be a useful palliative remedy for controlling seizures in patients with medication-resistant epilepsies.^4^ However, median seizure frequency reduction is only about two-thirds compared to baseline and few become seizure-free.^5,6^ This limitation underscores the need to deepen our understanding of how seizures originate and spread throughout the brain, as this knowledge provides the foundation for developing effective therapies.

As more data from thalamic recordings have emerged from two decades of pioneering work in epilepsy patients ^*7-15*^, our scientific understanding of seizure propagation through the human thalamus has been evolving but still remains significantly limited. Recent advances in multisite thalamic recordings in epilepsy patients have provided a new platform for recording across multiple sites of the thalamus simultaneously and mapping their involvement during seizure propagation in the human brain.

Prior intracranial recordings in the human thalamus suggested its involvement in majority of focal seizures, with early engagement observed not only in the anterior subregion of the thalamus (ANT) but also in the mediodorsal (MD) and pulvinar (PLV) subdivisions.^10^ To better understand these thalamic contributions, our previous work^14^ applied simultaneous recordings across multiple thalamic subdivisions in each patient, enabling direct comparisons of seizure involvement across specific thalamic regions (ANT, MD, and PLV subdivisions) within individual brains. Our analyses revealed that the medial PLV was more prominently engaged than was ANT during seizures—even in those originating from the medial temporal structures such as the hippocampus and amygdala. More importantly, we observed that seizures sharing the same onset signature (i.e., onset zone and electrographic features) exhibited a stereotyped pattern of thalamic engagement. In contrast, seizures with distinct onset patterns showed correspondingly different thalamic recruitment profiles. This suggests that variations in the thalamic footprint may reflect differences in the origin of the seizures and in the specific ictal networks involved. In our previous study, the role of MD in seizure propagation remained poorly characterized due to limited sampling. Indeed, the role of the MD subdivision of the thalamus in seizure propagation has remained insufficiently understood due to under sampling of this thalamic subregion in human studies. ^16^

Understanding the patterns and circuits that enable seizure propagation is not only a matter of scientific curiosity but also of significant clinical relevance. While achieving seizure freedom remains our ultimate goal, preventing seizures from spreading widely—and thereby causing potentially injurious or disabling symptoms—could in itself represent a major step forward in improving patient’s autonomy and quality of life. This underscores the critical need to characterize the specific pathways involved in seizure propagation.

In this context, MD thalamus becomes particularly intriguing. In most individuals, both thalami are interconnected through a midline structure known as the *massa intermedia (MI)*. Although some authors have suggested that this commissure lacks functional significance,^17^ growing evidence points toward potential inter-thalamic communication through the MI structure.^18,19^ This structural connection between the two thalami could be one possible path (among others) for seizure propagation from one hemisphere to the other—and thus a potentially promising target for neuromodulation aimed at preventing secondary generalization of seizures.

Building on this rationale, we conducted a stereoelectroencephalography (sEEG) study in 23 patients with multisite thalamic recordings (i.e., simultaneous and bilateral recordings of the ANT, MD, and PLV subdivisions). We used electrode trajectories traversing the MI to clinically assess bilateral involvement and this allowed assessment of inter-thalamic seizure propagation. In our analysis, we included several seizure types and onset zones extending beyond the temporal lobe. The study aimed to address three key questions: 1) Does the ictal activity in seizure onset zone involve the MD subregion of the thalamus? 2) Is MD involvement preferentially associated with specific seizure onset zones? And 3) Does the involvement of the MD correlate with propagation of ictal activity to the contralateral hemisphere?

## Materials and Methods

### Patients

From a larger cohort of patients who underwent thalamic implantation at Stanford Health Care between mid-2021 and 2025, we selected 23 patients for analysis. Inclusion required simultaneous recordings from the ipsilateral ANT, PLV, and MD sites. All patients presented with focal epilepsy of unclear lateralization or origin and were being considered for neuromodulatory therapy, particularly DBS targeting ANT, based on promising outcomes from the SANTE trial and subsequent studies.^4-6^ The details of the patients’ demographic data can be found in the Supplementary Materials (**Table S1**).

### Ethical Approval and Informed Consent

Prior studies have demonstrated the safety of sEEG recordings within various thalamic sites.^8,10,20-22^ Our approach extended previous protocols by recording from multiple thalamic regions concurrently. This enabled examination of intra-thalamic propagation dynamics. All participants provided informed consent, and the study protocol was approved by the Stanford Institutional Review Board (IRB).

### Electrode Planning and Target Selection

Electrode placement was determined in weekly multidisciplinary epilepsy surgery conferences involving epileptologists, neurosurgeons, neuroradiologists, neuropsychologists, and neuropsychiatrists. Planning incorporated clinical history, neurological exams, neuropsychological evaluations, and multimodal imaging (MRI, PET, SPECT, and, when available, MEG or high-density EEG). Decisions on thalamic targeting, including MD subregion coverage, were made based on both diagnostic and therapeutic considerations.

### Surgical Procedure and Thalamic Implantation

Thalamic targets were accessed by extending trajectories already used for cortical recordings. No additional electrodes were required solely for thalamic sampling, and cortical recordings were not compromised. Regions sampled included temporal, insular, frontal opercular, parietal, and occipital cortices. Thalamic sites—especially MD, ANT, and PLV—were targeted using an orthogonal approach,^14^ with bilateral coverage when feasible. To minimize tissue disruption, electrodes with 0.86 mm diameters (Ad-Tech Medical, Oak Creek, WI) and 3-5 mm inter-contact. More details about targeting, and intraoperative workflow for thalamic implantations has been described elsewhere.^23^

### Postoperative Imaging and Electrode Localization

Post-implant CT imaging confirmed electrode placement and ruled out hemorrhage. Electrode positions were mapped in FreeSurfer space using the iELVis toolbox.^24^ 3D anatomical reconstruction was performed using recon-all (FreeSurfer v6.0.0).^25^ Image registration was achieved with FSL’s flirt function^26,27^ and FreeSurfer’s bbregister.^28^ Electrodes were manually labeled in BioImage Suite,^29^ and coordinates were defined in each patient’s native space.

### EEG Acquisition

Continuous EEG was recorded at 1000 Hz using Nihon-Kohden systems with video monitoring. Thalamic and cortical signals were visualized using a bipolar montage, with filters adjusted based on region (0.001–0.1 s time constant; 300 Hz low-pass filter; 10 µV sensitivity). Artifact-prone channels were excluded from analysis.

### Seizure Classification

Seizures in this study were classified according to the ILAE 2025^30^ updated nomenclature. Due to the lack of a standardized method to assess consciousness, we classify seizures as focal seizures (FS) for events that were focal without secondary generalization and with clinical manifestations; focal-to-bilateral tonic-clonic seizures (FBTC) the focal seizures that evolved into generalized tonic-clonic activity; and finally, electrographic seizures (ES) were labeled for events detected only on sEEG recordings without clear clinical manifestations.

### EEG Analysis of Ictal Propagation

Seizure onset zones and early thalamic recruitment for all captured seizures were initially reviewed by the inpatient clinical team. Analysis of each seizure was by at least two board-licensed or eligible electroencephalographers using the following criteria: rhythms evolving in frequency, morphology or spread; emergence of rhythmic spiking or sharp activity, low-voltage fast activity, high-frequency oscillations, burst suppression, or delta.^31^ Location and primacy of onset of the seizure was based on visual inspection of the sEEG channels. Two epilepsy fellows (blinded to each other’s interpretations and to clinical data) then retrospectively re-evaluated a subset of seizures (up to three seizures per seizure subtype for all patients) to determine onset patterns and thalamic propagation. A senior EEG board-licensed epileptologist resolved any discrepancies. Propagation patterns were assessed for thalamic engagement, subregion of earliest involvement (with emphasis on MD), and temporal sequencing of activity. Timing data were analyzed at the second level to avoid overstating precision. As noted later in the Discussion, we avoided computational methods to ensure practical feasibility and ecological validity. However, we acknowledge that the absence of unbiased computational approaches is a limitation of our study.

### Assessment of the Massa Intermedia

The presence and morphology of the massa intermedia were assessed through visual inspection of each patient’s structural MRI by a board-certified neuroradiologist and a radiology resident. The massa intermedia was considered present if a continuous gray matter bridge connecting the medial thalami was visible in coronal and axial planes.

### Statistical Analysis

In the current study, we mainly focus on providing descriptive statistics.

## Results

### Patient Demographics

23 patients (12 male, 11 female, 20-57 years old, 35.7 mean age) were included in this study. The sole inclusion criterion in the study was the presence of bilateral thalamic coverage in the mediodorsal region of the thalamus simultaneously with ANT and PLV subregions unilaterally or bilaterally. Demographic data can be found in the Supplementary Materials (**Table S1**).

### Electrode Coverage

In total, we had 103 electrodes in the thalamus including 40 anterior, 23 mid, and 40 posterior subdivisions (**Table S1**). In this study, thalamic recordings are reported using subregions, subdivisions, or thalamic sites rather than exact nuclei, reflecting the technical, anatomical, and atlas-related limitations of SEEG in humans. This approach allows us to apply a standardized framework across all patients and obtain reliable and reproducible results.

### Safety

The procedure was well tolerated by all patients, without any complications. Notably, there were no significant hemorrhages in the thalamus, except for one case involving a parietal hemorrhage in an extrathalamic electrode. All patients returned to their preoperative baseline following electrode implantation and were able to fully participate in the necessary clinical assessments. As expected, transient pneumocephalus was uniformly observed after electrode explantation, but no clinical complications were attributed to the removal of the electrodes.

### Seizure classification

A total of 119 seizures from 23 patients were reviewed. Our analysis comprised spontaneous seizures (i.e., not triggered by electrical stimulation). Seizure onset zones (SOZ) were identified based on visual inspection of sEEG recordings across all implanted brain regions. Seizures originated from the temporal lobe, orbitofrontal, frontal, parietal, occipital and cingulate regions. For patients whose seizures showed simultaneous onset in two regions, we created combined categories such as fronto-temporal for a patient with concurrent frontal and temporal onset, and temporo-occipital for a patient with simultaneous temporal and occipital involvement. Majority of seizures originated from the temporal lobe (75, 63.0%), followed by orbitofrontal (13, 10.9%), fronto-temporal (9, 7.6%), occipital (9, 7.6%), frontal (5, 4.2%), parietal (4, 3.4%), cingulate (2, 1.7%), and temporo-occipital (2, 1.7%) regions.

### Clinical seizure types

In our cohort, seizures were mainly focal seizures without secondary generalization (FS, 90 seizures in 21 patients), but we also captured data from focal-to-bilateral tonic–clonic seizures (FBTC, 11 seizures in nine patients), and electrographic seizures (ES, 18 seizures in six patients).

### Thalamic involvement at seizure onset

Distinct activation patterns emerged across the 119 selected seizures. **Table 1** provides details of all seizures.

**Table 1.**
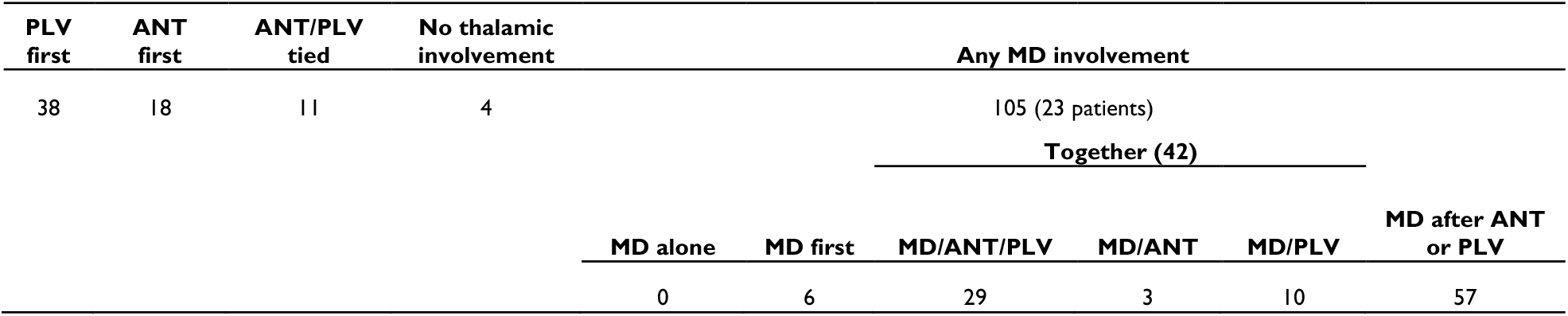
Thalamic subregions at seizure onset. Overview of thalamic subregions recruited at seizure onset, showing the frequency of involvement of the MD, ANT, and PLV sites. The table specifies whether each thalamic site was involved alone, in combination, or simultaneously with others. Within the cases involving the MD subregion, a further subclassification was performed according to the temporal sequence of recruitment and co-involvement with other thalamic sites.

#### Thalamic Involvement during Seizures

In 96.6% of seizures, our visual inspection of the EEGs recorded from the thalamus showed clear ictal changes in the thalamic recordings. In three patients with bilateral multisite sampling of the thalamus, we could not detect visual signs of thalamic involvement in four (3.4%) seizures – all of which originated in the temporal lobes. However, in the same patients, other seizures were recorded with clear thalamic engagement. None of the four seizures had signs of propagation to the contralateral hemisphere.

#### Timing of Thalamic Involvement

Among the 115 seizures with thalamic involvement, 99 (86%) showed thalamic involvement within the first 15 seconds of seizure onset. Notably, all FBTC seizures exhibited very rapid thalamic involvement, with thalamic recruitment occurring within the first 5 seconds in eight out of 11 cases (72.7%). Similarly, early thalamic engagement (≤15 s) was observed in 77 of 90 FS seizures (85.5%) and in 11 of 18 ES seizures (61%).

#### Thalamic Site First Involved

The PLV subregion was the first thalamic structure to be activated in 38 seizures (31.9%), majority of which originated from the temporal lobe. The ANT was first involved in 18 seizures (15.1%), of which only seven had a temporal onset. The MD site was the first thalamic site to be recruited in only 6 seizures (5%), all of which originated from the temporal lobes (five of six from the MTL temporal lobes).

#### Multiple Thalamic Sites First Involved

29 seizures (24.4%) showed apparently simultaneous involvement of all three thalamic sites (ANT/PLV/MD) during seizure propagation, indicating coordinated thalamic engagement rather than sequential recruitment. In the remaining cases, either one or a pair of thalamic sites were engaged (see **Table 1** for details).

#### MD Involvement during Seizures

The MD site was involved in 105 of 119 seizures (88.2%) overall. However, it was never the *only* thalamic subregion to be engaged without the other thalamic sites also being engaged. Comparing the temporal order of MD and other thalamic sites’ involvement, we found that the MD showed involvement before or simultaneously with the other thalamic sites in 48 (40%) (See **Table 1** for details). In contrast, in 57 seizures, the MD was recruited after activation of the ANT and/or PLV, suggesting that, in majority of cases, the MD follows the initial activation of the other thalamic sites.

#### Thalamic Involvement and Seizure Onset Zones

Focusing on the data from patients with temporal lobe seizures (n = 75), we found that the PLV was involved first in 34.7% of cases, the ANT in 9.3%, and the MD in 8% of seizures. Simultaneous activation of all three sites (ANT/PLV/MD) was observed only in 16% of seizures. **Table 2** provides more details. Seizures engaging the MD subregion (105 of 119; **Table 3**), the temporal lobe was the most predominant source of seizures (66/105), of which 32 were from the mesial structures and 34 from neocortical temporal lobe regions. Majority of seizures with MD involvement showed spread to the contralateral hemisphere (**Table 3)**.

**Table 2.**
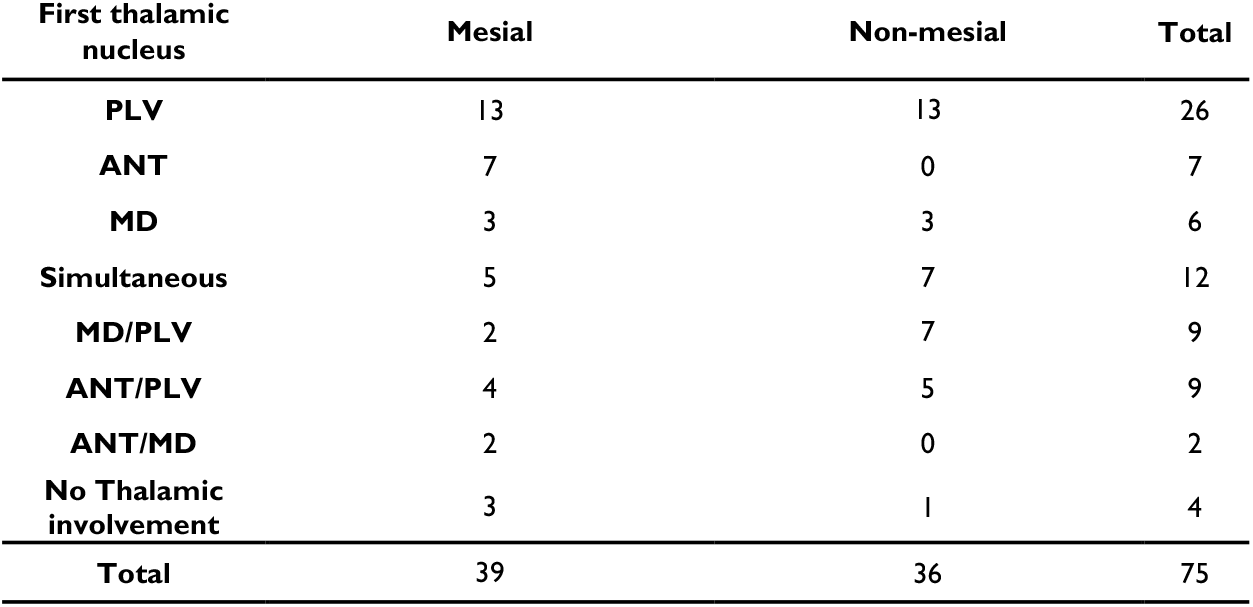
Temporal lobe seizures categorized by first thalamic subregion involvement and mesial vs. non-mesial onset. Distribution of temporal lobe seizures according to the first thalamic nucleus involved and the site of cortical onset (strictly mesial vs. non-mesial or broader onset).

| First thalamic nucleus | Mesial | Non-mesial | Total |
| --- | --- | --- | --- |
| PLV | 13 | 13 | 26 |
| ANT | 7 | 0 | 7 |
| MD | 3 | 3 | 6 |
| Simultaneous | 5 | 7 | 12 |
| MD/PLV | 2 | 7 | 9 |
| ANT/PLV | 4 | 5 | 9 |
| ANT/MD | 2 | 0 | 2 |
| No Thalamic involvement | 3 | 1 | 4 |
| Total | 39 | 36 | 75 |

**Table 3.** Distribution of seizure onset regions according to MD involvement. Numbers indicate the absolute count of seizures in each onset region, with the numbers in parentheses representing seizures that did exhibit contralateral spread. Temporal lobe seizures are further subdivided into Mesial only (restricted to amygdala/hippocampus) and Broader/Other (mesial with broader onset/lateral, inferior, or superior temporal regions). MD involvement refers to whether the mediodorsal subregion was involved in the seizure (Yes) or not (No).

| MD involvement | Orbitofrontal | Frontal | Parietal | Cingulate | Temporal |  | Occipital | Temporo-occipital | Fronto-Temporal |
| --- | --- | --- | --- | --- | --- | --- | --- | --- | --- |
|  |  |  |  |  | Strictly Mesial | Other in temporal |  |  |  |
| Yes (n=105) | 12 | 5 | 4 | 1 | 32 (21) | 34 (21) | 9 | 2 | 6 |
| No (n=14) | 1 | 0 | 0 | 1 | 7 (0) | 2 (1) | 0 | 0 | 3 |

### Contralateral spread and MD involvement

In 78 (65.5%) seizures, ictal activity reached one or more recording sites in the contralateral hemisphere during seizure propagation. The MD site ipsilateral to the seizure onset side was involved 96.2% of these seizures while the contralateral MD site was involved in 94.9% (See more details in **Table 4a and 4b**). Interestingly, in 14 seizures where we could not identify visual signs of seizure engagement in the ipsilateral MD, only three had signs of ictal spread to any of the contralateral recording sides.

**Table 4.** MD involvement in seizures with and without contralateral spread. In this table we describe Ipsilateral MD (4a) and contralateral MD (4b) involvement seizures with and without contralateral spread. Values are presented in the format X (Y), where X represents the number of seizures and Y.

| Ipsilateral Seizure Involvement |  |  |  |  | Contralateral Seizure Involvement |  |  |  |  |
| --- | --- | --- | --- | --- | --- | --- | --- | --- | --- |
| Ipsi MD |  | Yes | No | Total | CTL MD |  | Yes | No | Total |
| Yes | Pre-Gz | 51 (18) | 30 (9) | 105 | Yes | Pre-Gz | 47 (18) | 25 (9) | 99 |
|  | Post-Gz | 24 (10) |  |  |  | Post-Gz | 27 (12) |  |  |
| No |  | 3 (2) | 11 (7) | 14 | No |  | 4 (3) | 16 (9) | 20 |
| Total |  | 78 | 41 | 119 | Total |  | 78 | 41 | 119 |

Among the 78 seizures with contralateral propagation, 28 (36%) followed the pathway of ipsilateral MD to contralateral MD and then to contralateral cortex; 19 (24%) propagated simultaneously to the ipsilateral MD and the contralateral cortex; 23 (30%) spread to the contralateral cortex first before contralateral thalamic involvement; and 3 (4%) propagated to the mPLV and/or ANT subregions before reaching the contralateral cortex. The remaining five seizures could not be classified due to either lack of contralateral thalamic sampling (two seizures) or absence of thalamic involvement (three seizures; **Table 5**).

**Table 5.** First Site of Contralateral Involvement: Cortex vs. ANT/PLV. Distribution of first sites of contralateral involvement across seizures, comparing early recruitment of the contralateral cortex versus ANT and or PLV thalamic subregions.

| Contralateral nuclei first (ANT and/or PLV) | Cortex first |  |  | Simultaneous Cortex with ANT and/or PLV | No contralateral nuclei sampled |
| --- | --- | --- | --- | --- | --- |
| 3 | 19 |  |  | 4 | 2 |
|  | Then ANT | Then PLV | Then ANT/PLV |  |  |
|  | 8 | 6 | 5 |  |  |

It is noteworthy that in majority (61%) of the seizures with contralateral spread we could observe signs of ictal involvement in the contralateral MD prior to or simultaneous with ictal spread to other contralateral recording sites.

When analyzing the corresponding seizure onset zones, seizures that propagated first to the MD were predominantly of mesial temporal (MTL) origin (57%), whereas, if the contralateral cortex was involved before the contralateral MD, 89% had neocortical (non-MTL) onset. More details are presented in **Table 6**.

**Table 6.** Distribution of seizure onset locations by first activated structure during contralateral spread. This table shows the distribution of seizure onset locations among seizures that exhibited contralateral spread. Seizures are classified according to whether the first contralateral structure involved was the contralateral MD, the contralateral cortex or both synchronous. Temporal lobe onsets are further subdivided into *mesial* and *lateral/other* temporal region.

| First activated structure | Temporal |  | Orbitofrontal | Frontal | Parietal | Fronto-temporal | Occipital | Temporo-occipital | Cingulate | Total seizures |
| --- | --- | --- | --- | --- | --- | --- | --- | --- | --- | --- |
|  | Mesial | Lateral/other |  |  |  |  |  |  |  |  |
| Contralateral MD | 16 | 5 | 3 | 1 | 1 | 0 | 0 | 2 | 0 | 28 |
| Contralateral Cortex | 2 | 7 | 1 | 0 | 1 | 1 | 7 | 0 | 0 | 19 |
| Synchronous MD-Cortex | 3 | 5 | 4 | 1 | 3 | 2 | 0 | 0 | 1 | 19 |

## Discussion

### Thalamic involvement is observed in all patients

In our cohort, every patient presented seizures with clear thalamic recruitment. Early thalamic engagement (≤15 s from seizure onset) was observed in 83% of seizures overall, and in 73% of focal-to-bilateral tonic-clonic seizures, the thalamus was recruited within the first 5 seconds. These findings align with prior studies reporting rapid thalamic recruitment in temporal lobe epilepsy,^8,15^ with up to 86% of seizures showing thalamic involvement within the first 15 seconds. They also reported that 80% of mesial temporal lobe epilepsy cases had thalamic involvement. Our findings extend these observations, showing even more prominent involvement: 95% of the temporal seizures and 92% of the MTL seizures exhibited thalamus involvement.

### The MD subregion is rarely the first thalamic site recruited but is highly involved in MTL seizures

The MD was the *first* thalamic site in only 5% of seizures, all originating in temporal lobe regions and the majority were from MTL. Importantly, the MD was never recruited in isolation, always co-activated with the other thalamic sites. It was involved in every patient and was engaged in 88% of the seizures. Looking specifically at MTL seizures, the MD was involved in 82% and very interestingly, in the rare MTL seizures where the MD was not recruited, contralateral propagation never occurred.

Rodent studies further supported this MD-MTL connection, showing that MD stimulation in piriform or hippocampal kindled rats reduced seizure duration by 80% and sometimes suppressed seizures, observing that the most effective region was dorsolateral at the axon entry zone. ^32^ These results are also in line with a new connectivity mapping study ^33^ in the human brain reporting that the MD thalamic subregion received disproportionately strong projections from the MTL than it sent to MTL (stronger MD inflow from MTL), highlighting markedly asymmetric connectivity between the MD and MTL regions, while the MD sends strong projections to the cingulate, insular, and prefrontal cortices.

**We replicated our prior findings from the ANT and PLV**, which reported that the PLV subregion was engaged earlier and more prominently than was the ANT in a majority of patients.^14^ In our current analysis the PLV was the first thalamic subregion recruited in a substantial proportion of temporal epilepsy cases (35%), while the ANT was the first thalamic structure just in 9% of the temporal seizures. In previous reports, McGinn et al. ^15^ also showed in a direct comparison of simultaneous ANT and PLV recordings that PLV site was activated first in 82% of patients, whereas initial ANT activation occurred in only ∼31%.^15^ These findings are consistent with the hypothesis that the PLV subregion contributes earlier and more prominently to temporal lobe seizure propagation compared with the ANT or MD.

### The MD may play a key role in contralateral seizure propagation, particularly from mesial temporal onset zones

Among seizures with interhemispheric spread, the ipsilateral MD was engaged in 96.2% and the contralateral MD in 94.9%. In 61% of the seizures with contralateral propagation, the contralateral MD was recruited prior to or simultaneously with the contralateral cortex, suggesting a potential role in seizure propagation to the contralateral hemisphere. Seizures propagating *first* through the MD were predominantly of MTL origin, whereas those involving the contralateral cortex before the contralateral MD mostly had neocortical onset and a more heterogeneous distribution (Orbitofrontal, Frontoparietal, Temporal, Parietal, Occipital). These results may indicate that both thalamo-cortical and cortico-cortical pathways contribute to interhemispheric propagation. While the corpus callosum has traditionally been considered the primary conduit for seizures crossing hemispheres, and callosotomy is used to treat pharmaco-resistant epilepsy with rapidly generalizing or drop attacks, approximately 80% of patients do not achieve complete seizure freedom, and nearly half continue to experience drop attacks post-surgery.^34^ This clinical evidence highlights that additional pathways, including thalamic circuits such as the MD, may contribute to interhemispheric seizure propagation.

### Evidence from human imaging and electrophysiology supports the translational relevance of the MD subregion in seizure propagation, although data remain limited

Keller et al^35^ reported thalamo-temporal atrophy in temporal lobe epilepsy patients. Subsequent studies indicated that surgical non-responders tended to exhibit bilateral atrophy of the MD and pulvinar subregions.^35,36^ Neurophysiological studies using deep brain stimulation of thalamic subregions have further demonstrated that stimulating the MD subdivision predominantly activated the ipsilateral orbitofrontal cortex and mesial and lateral frontal regions, but also recruited mesial temporal structures.^37^ It is also known that the MD is implicated in cognitive functions such as working memory and executive control, which may transiently be affected during seizures, highlighting the functional importance of this subregion beyond seizure propagation.^38,39^

### Clinical Implications

Taken together, our findings may outline a model of seizure propagation-especially for those originating in the MTL-traveling through thalamic structures via the MD subregion and massa intermedia and subsequently reaching other brain regions where outflow interactions are strongest. Its consistent recruitment during seizure spread makes this thalamic subregion a potential target for neuromodulatory therapies though caution is warranted given potential cognitive effects^39^ and reports of unpleasant subjective experiences during stimulation^33^.

Future studies are needed to learn how thalamic and interhemispheric networks shape seizure propagation, but each insight we gain brings us a step closer to optimize therapies that can better control seizures and improve daily life. While epilepsy remains a complex challenge, our ultimate goal is clear: to translate this knowledge into meaningful benefits for the patients we care for.

### Our study has its limitations

A key limitation of our study is the high proportion of temporal lobe seizures, reflecting the higher prevalence of temporal lobe epilepsies (TLE) in the general population. This imbalance may bias our findings toward TLE, and thus some observations may not generalize to all epilepsy types. Additionally, we were unable to perform a detailed analysis of awareness during seizures. As a result, all focal events were classified as focal seizures without generalization, without determining patient’s level of consciousness, limiting our ability to explore the relationship between thalamic engagement and level of consciousness during seizures. Furthermore, we did not use computational tools for the analysis of thalamic engagement during seizures because such tools are only available in laboratories and have not been integrated to routine clinical practice. This was clearly a limitation of our study compared to several other important studies of the thalamus in the human brain^8-10,40-43^. Our rationale for avoiding computational methods was to ensure that our results are practically feasible and ecologically valid, but we are mindful that we cannot assess to what extent such observer-dependent methods are affected by the observer’s inexperience or bias. We are hopeful that objective computational tools will eventually become widely available in our everyday clinical practice of epileptology and will be taken into use to allay concerns about reviewer inexperience or bias.

## References

1. Davis P, Gaitanis J. Neuromodulation for the Treatment of Epilepsy: A Review of Current Approaches and Future Directions. Clin Ther. Jul 2020;42(7):1140–1154. doi:10.1016/j.clinthera.2020.05.017

2. Li MCH, Cook MJ. Deep brain stimulation for drug-resistant epilepsy. Epilepsia. Feb 2018;59(2):273–290. doi:10.1111/epi.13964

3. Salanova V. Deep brain stimulation for epilepsy. Epilepsy Behav. Nov 2018;88S:21–24. doi:10.1016/j.yebeh.2018.06.041

4. Fisher R, Salanova V, Witt T, et al. Electrical stimulation of the anterior nucleus of thalamus for treatment of refractory epilepsy. Epilepsia. May 2010;51(5):899–908. doi:10.1111/j.1528-1167.2010.02536.x

5. Salanova V, Witt T, Worth R, et al. Long-term efficacy and safety of thalamic stimulation for drug-resistant partial epilepsy. Neurology. Mar 10 2015;84(10):1017–25. doi:10.1212/WNL.0000000000001334

6. Salanova V, Sperling MR, Gross RE, et al. The SANTE study at 10 years of follow-up: Effectiveness, safety, and sudden unexpected death in epilepsy. Epilepsia. Jun 2021;62(6):1306– 1317. doi:10.1111/epi.16895

7. Romeo A, Issa Roach AT, Toth E, et al. Early ictal recruitment of midline thalamus in mesial temporal lobe epilepsy. Ann Clin Transl Neurol. Aug 2019;6(8):1552–1558. doi:10.1002/acn3.50835

8. Guye M, Regis J, Tamura M, et al. The role of corticothalamic coupling in human temporal lobe epilepsy. Brain. Jul 2006;129(Pt 7):1917–28. doi:10.1093/brain/awl151

9. Chaitanya G, Toth E, Pizarro D, Irannejad A, Riley K, Pati S. Precision mapping of the epileptogenic network with low- and high-frequency stimulation of anterior nucleus of thalamus. Clin Neurophysiol. Sep 2020;131(9):2158–2167. doi:10.1016/j.clinph.2020.05.036

10. Pizzo F, Roehri N, Giusiano B, et al. The Ictal Signature of Thalamus and Basal Ganglia in Focal Epilepsy: A SEEG Study. Neurology. Jan 12 2021;96(2):e280–e293. doi:10.1212/WNL.0000000000011003

11. Gadot R, Korst G, Shofty B, Gavvala JR, Sheth SA. Thalamic stereoelectroencephalography in epilepsy surgery: a scoping literature review. J Neurosurg. Mar 11 2022:1–16. doi:10.3171/2022.1.JNS212613

12. Piper RJ, Richardson RM, Worrell G, et al. Towards network-guided neuromodulation for epilepsy. Brain. Jun 30 2022; doi:10.1093/brain/awac234

13. Ilyas A, Tandon N, Lhatoo SD. Thalamic neuromodulation for epilepsy: A clinical perspective. Epilepsy Res. Jul 2022;183:106942. doi:10.1016/j.eplepsyres.2022.106942

14. Wu TQ, Kaboodvand N, McGinn RJ, et al. Multisite thalamic recordings to characterize seizure propagation in the human brain. Brain. Jul 3 2023;146(7):2792–2802. doi:10.1093/brain/awad121

15. McGinn R, Von Stein EL, Datta A, et al. Ictal Involvement of the Pulvinar and the Anterior Nucleus of the Thalamus in Patients With Refractory Epilepsy. Neurology. Dec 10 2024;103(11):e210039. doi:10.1212/WNL.0000000000210039

16. Salami P, Paulk AC, Soper DJ, et al. Inter-seizure variability in thalamic recruitment and its implications for precision thalamic neuromodulation. Commun Med (Lond). May 22 2025;5(1):190. doi:10.1038/s43856-025-00920-9

17. Parra JED, Ripoll AP, Garcia JFV. Interthalamic adhesion in humans: a gray commissure? Anat Cell Biol. Mar 31 2022;55(1):109–112. doi:10.5115/acb.21.164

18. Sahin MH, Gungor A, Demirtas OK, et al. Microsurgical and fiber tract anatomy of the interthalamic adhesion. J Neurosurg. Nov 1 2023;139(5):1386–1395. doi:10.3171/2023.3.JNS221669

19. Borghei A, Kapucu I, Dawe R, Kocak M, Sani S. Structural connectivity of the human massa intermedia: A probabilistic tractography study. Hum Brain Mapp. Apr 15 2021;42(6):1794–1804. doi:10.1002/hbm.25329

20. Chaitanya G, Romeo AK, Ilyas A, et al. Robot-assisted stereoelectroencephalography exploration of the limbic thalamus in human focal epilepsy: implantation technique and complications in the first 24 patients. Neurosurg Focus. Apr 1 2020;48(4):E2. doi:10.3171/2020.1.FOCUS19887

21. Arthuis M, Valton L, Regis J, et al. Impaired consciousness during temporal lobe seizures is related to increased long-distance cortical-subcortical synchronization. Brain. Aug 2009;132(Pt 8):2091–101. doi:10.1093/brain/awp086

22. Gadot R, Korst G, Shofty B, Gavvala JR, Sheth SA. Thalamic stereoelectroencephalography in epilepsy surgery: a scoping literature review. J Neurosurg. Nov 1 2022;137(5):1210–1225. doi:10.3171/2022.1.JNS212613

23. Jamiolkowski RM, Datta A, Willsey MS, Parvizi J, Buch VP. Multinuclear thalamic targeting with human stereotactic electroencephalography: surgical technique and nuances. J Neurosurg. Apr 1 2025;142(4):936–944. doi:10.3171/2024.7.JNS24452

24. Groppe DM, Bickel S, Dykstra AR, et al. iELVis: An open source MATLAB toolbox for localizing and visualizing human intracranial electrode data. J Neurosci Methods. Apr 01 2017;281:40–48. doi:10.1016/j.jneumeth.2017.01.022

25. Fischl B. FreeSurfer. Neuroimage. Aug 15 2012;62(2):774–81. doi:10.1016/j.neuroimage.2012.01.021

26. Jenkinson M, Beckmann CF, Behrens TE, Woolrich MW, Smith SM. Fsl. NeuroImage. Aug 15 2012;62(2):782–90. doi:10.1016/j.neuroimage.2011.09.015

27. Jenkinson M, Smith S. A global optimisation method for robust affine registration of brain images. Med Image Anal. Jun 2001;5(2):143–56.

28. Greve DN, Fischl B. Accurate and robust brain image alignment using boundary-based registration. NeuroImage. Oct 15 2009;48(1):63–72. doi:10.1016/j.neuroimage.2009.06.060

29. Papademetris X, Jackowski MP, Rajeevan N, et al. BioImage Suite: An integrated medical image analysis suite: An update. Insight J. 2006;2006:209.

30. Ragunathan K, Veeraraghavan V, Kaushik JS. ILAE 2025 Classification of Epileptic Seizures: Key Revisions and Implications for Clinical Practice. Indian Pediatr. Aug 2025;62(8):623–627. doi:10.1007/s13312-025-00120-7

31. Perucca P, Dubeau F, Gotman J. Intracranial electroencephalographic seizure-onset patterns: effect of underlying pathology. Brain. Jan 2014;137(Pt 1):183–96. doi:10.1093/brain/awt299

32. Zhang DX, Bertram EH. Suppressing limbic seizures by stimulating medial dorsal thalamic nucleus: factors for efficacy. Epilepsia. Mar 2015;56(3):479–88. doi:10.1111/epi.12916

33. Pantis S, Lyu D, Quabs J, et al. Electrophysiological Brain Connectivity and Subjective States Evoked by Electrical Stimulation of the Human Mediodorsal Thalamus. bioRxiv. 2025:2025.11.12.688132. doi:10.1101/2025.11.12.688132

34. Chan AY, Rolston JD, Lee B, Vadera S, Englot DJ. Rates and predictors of seizure outcome after corpus callosotomy for drug-resistant epilepsy: a meta-analysis. J Neurosurg. Apr 1 2019;130(4):1193–1202. doi:10.3171/2017.12.JNS172331

35. Keller SS, O’Muircheartaigh J, Traynor C, Towgood K, Barker GJ, Richardson MP. Thalamotemporal impairment in temporal lobe epilepsy: a combined MRI analysis of structure, integrity, and connectivity. Epilepsia. Feb 2014;55(2):306–15. doi:10.1111/epi.12520

36. Keller SS, Richardson MP, Schoene-Bake JC, et al. Thalamotemporal alteration and postoperative seizures in temporal lobe epilepsy. Ann Neurol. May 2015;77(5):760–74. doi:10.1002/ana.24376

37. Zumsteg D, Lozano AM, Wieser HG, Wennberg RA. Cortical activation with deep brain stimulation of the anterior thalamus for epilepsy. Clin Neurophysiol. Jan 2006;117(1):192–207. doi:10.1016/j.clinph.2005.09.015

38. Golden EC, Graff-Radford J, Jones DT, Benarroch EE. Mediodorsal nucleus and its multiple cognitive functions. Neurology. Nov 15 2016;87(20):2161–2168. doi:10.1212/WNL.0000000000003344

39. Peräkylä J, Sun L, Lehtimäki K, et al. Causal evidence from humans for the role of mediodorsal nucleus of the thalamus in working memory. J Cogn Neurosci. Dec 2017;29(12):2090–2102. doi:10.1162/jocn_a_01176

40. Soulier H, Pizzo F, Jegou A, et al. The anterior and pulvinar thalamic nuclei interactions in mesial temporal lobe seizure networks. Clin Neurophysiol. Jun 2023;150:176–183. doi:10.1016/j.clinph.2023.03.016

41. Evangelista E, Benar C, Bonini F, et al. Does the Thalamo-Cortical Synchrony Play a Role in Seizure Termination? Front Neurol. 2015;6:192. doi:10.3389/fneur.2015.00192

42. Bartolomei F, Chauvel P, Wendling F. Epileptogenicity of brain structures in human temporal lobe epilepsy: a quantified study from intracerebral EEG. Brain. Jul 2008;131(Pt 7):1818–30. doi:10.1093/brain/awn111

43. Ilyas A, Toth E, Chaitanya G, Riley K, Pati S. Ictal high-frequency activity in limbic thalamic nuclei varies with electrographic seizure-onset patterns in temporal lobe epilepsy. Clin Neurophysiol. May 2022;137:183–192. doi:10.1016/j.clinph.2022.01.134

