## Supplementary Material for "Human Mediodorsal Thalamus in Seizure Propagation"

**Table S1. Demographics**

| Patient Number | Sex | Age | Handedness | Age of onset | Imaging Findings (MRI/FDG-PET) | Seizure Frequency (seizures/month) | Anti-seizure medications | Seizure focus | Number of depth electrodes | Number of contacts | Thalamic implantation |
|---|---|---|---|---|---|---|---|---|---|---|---|
| 2 | M | 40 | RHD | 16 | Globular L hippocampus; subtle L mesial temporal hypometabolism | FIAS several per week; FTBTC 2-3/ month | Onfi 5 mg daily, Trileptal 600 mg BID, Fycompa 6 mg daily | L fusiform gyrus | 10 | 116 | Bilateral MD, only L ANT & L PLV |
| 3 | M | 52 | RHD | 17 | Non-lesional; slight right frontal and right anterior temporal hypometabolism. | FIAS 2-3/week, rare FTBTC | Lamotrigine 150 mg BID, Keppra 1000 mg BID, Dilantin 300 mg qHS | (1) L lateral temporal, (2) R orbitofrontal | 21 | 252 | Bilateral ANT, PLV, and MD |
| 4 | M | 28 | RHD | 20 | Small left anterior ventral temporal lobe dysplasia; decreased uptake in the right temporal lobe and left inferior temporal gyrus. | FAS 2/month. FIAS, FTBTC with variable frequency | Levetiracetam 2000 mg BID, Lacosamide 200 mg BID | 1) Left anterior & inferior temporal lobe - where dysplasia was newly discovered, 2) R hippocampus & amygdala | 20 | 252 | Bilateral ANT, PLV, and MD |
| 5 | F | 23 | RHD | 18 | Globular left hippocampus with incomplete hippocampal inversion; mildly decreased uptake within the left mesial temporal lobe. | FIAS, FTBTC 2/month | Lacosamide 200 mg BID | R inferior/mesial occipital region | 17 --> 25 | 228 --> 254 | Bilateral ANT, PLV, and MD |
| 6 | F | 37 | RHD | 10 | Non-lesional?; hypometabolism L>R mesial temporal regions | GTC 1/month, FIAS rarely, FAS 2x/week | Briviact 100 mg BID, lamotirigne 200 mg BID, Zonisamide 200 mg BID | L>R frontotemporal | 22 | 260 | Bilateral ANT, PLV, and MD |
| 7 | F | 35 | RHD | 15 | Prior left temporal lobectomy with stable postoperative changes; decreased uptake of L lateral temporal lobe | FIAS 2/month, GTC 1-2/month | Vimpat 200 BID, Lamotrigine 300 BID, Keppra XR 1500 BID | 1) Right posterior hippocampus & amygdala, 2) Left inferior temporal | 16 | 206 | Bilateral ANT, PLV, and MD |
| 8 | F | 47 | RHD | 30 | Mildly increased T2 flair signal intensity of left hippocampus & throughout L temporal lobe; mildly decreased FDG in left temporal lobe & insula | FIAS 3-5/week, single lifetime nocturnal convulsion | Lamotrigine 200 mg BID | 1) Left temporal lobe & insula, 2) Right temporal lobe & insula | 11 | 138 | Bilateral MD, only L ANT & L PLV |
| 9 | F | 40 | RHD | 23 | Left > right anterior temporal lobe encephalomalacia without significant hippocampal change; decreased uptake in left temporal pole | FIAS monthly, FAS 3-10/day, GTC not for many years | Lamotrigine 200 mg qAM & 300 mg qPM, Clonazepam 0.5 mg qPM, Eslicarbazepine 1200 mg qHS, VNS | Left orbitofrontal | 12 | 146 | Bilateral MD, bilateral ANT, only L PLV |
| 10 | M | 20 | RHD | 5 | Subtle abnormality of left inferior frontal region and minimal R hippocampal volume loss; subtle R>L mesial temporal hypometabolism | FIAS 2-3/day, FIAS 2-3/ year | Keppra XR 1000 mg qHS, Lamotrigine ER 200 mg qHS, Valproic acid 1250 mg qHS | Right SMA | 16 --> 21 | 212 | Bilateral ANT, PLV, and MD |
| 11 | M | 38 | RHD | 35 | Severe bilateral MTS and moderate diffuse volume loss; severe symmetric hypometabolism of bilateral anterior temporal lobes and bilateral post-central gyri | FIAS 3-4/week, GTC 3 lifetime episode | Vimpat 200 mg BID, VPA 500 mg BID, Zonisamide 200 mg qHS | 1) Left amygdala/hippocampus, 2) Right amygdala/hippocampus | 10 | 188 | Bilateral ANT, PLV, and MD |

| | | | | | | | | | | | |
|---|---|---|---|---|---|---|---|---|---|---|---|
| 12 | M | 36 | RHD | 16 | Small focal T2/FLAIR white matter signal abnormality in the right inferior anterior frontal lobe; normal FDG-PET | FAS 1-5/year | Vimpat 250 mg BID | Right superior temporal gyrus | 10 | 178 | Bilateral ANT, PLV, and MD |
| 13 | M | 51 | RHD | 28 | Left frontal lobe encephalomalacia; no PET available | FIAS/FTBTC 3-4/month | Vimpat 200 mg BID, Divalproex ER 500 mg qAM & 750 mg qHS | Left amygdala/hippocampus | 18 | 232 | Bilateral MD, only L ANT & L PLV |
| 14 | M | 57 | RHD | 30 | Single focus of T2 prolongation in the left posterior temporal lobe | FIAS 1/week, FAS 2-3/week | Keppra ER 1500 mg BID, lamotrigine ER 225 mg BID | Multifocal | 19 | 250 | Bilateral ANT, PLV, and MD |
| 16 | F | 32 | RHD | 19 | Non-lesional; subtle hypometabolism in the anterolateral right temporal lobe | FAS several per week, FIAS 1x/week, FTBTC 1-2x/month | Keppra 2000 mg BID, Trileptal 750 mg TID | Right posterior superior temporal gyrus | 15 | 198 | Bilateral ANT, PLV, and MD |
| 18 | M | 20 | RHD | 18 | Transient enlargement of the left temporal lobe that resolved on follow up imaging; FDG-PET normal | FIAS 1-3/month, FTBTC 3 lifetime events | LTG ER 200 BID, Vimpat 100 BID, Keppra XR 1000 BID, Lyrica 100 BID | 1) Bilateral anterior temporal lobes; 2) Left posterior temporal acute reactive seizure post-op thought likely due to left posterior hematoma | 17 | 216 | Bilateral ANT, PLV, and MD |
| 19 | F | 28 | RHD | 13 | L>R heterotopic gray matter along occipital & temporal horns of lateral ventricles & focal cortical dysplasia of right occipital lobe; diffusely decreased uptake of the left hemisphere in comparison to right | FIAS 1-3/month, FTBTC 1/year | Keppra 650 mg TID, Carbamazepine XR 400 mg TID | Left amygdala | 17 | 184 | Bilateral MD, bilateral PLV, L ANT |
| 20 | F | 27 | RHD | 14 | MRI normal, FDG-PET normal | Gelastic FIAS 1-3/month, FTBTC 1-2/month | Lamotrigine XR 200 mg daily, XCopri 200 mg daily | Multifocal | 18 | 232 | Bilateral ANT, PLV, and MD |
| 22 | M | 31 | RHD | 20 | Non-lesional; Minimal hypometabolism in the right anterior temporal and mesiotemporal regions | FAS 2/week, FTBTC once/month | Briviact 100 mg BID, Onfi 10 mg qAM & 15 mg qPM, Vimpat 200 mg BID | Bilateral hippocampi | 13 | 152 | Bilateral ANT, PLV, and MD |
| 23 | M | 27 | LHD | 22 | Non-lesional; left anterior temporal hypometabolism | FIAS every week, FTBTC less than once per month | Vimpat 300 mg BID, Clonazepam 2 mg daily | Left amygdala | 14 | 200 | Bilateral MD, bilateral ANT, L PLV |
| 24 | M | 29 | RHD | 16 | Post-surgical changes of LITT to right parietal/right occipital horn gray matter heterotopia & mild increased FLAIR signal and decreased volume of R hippocampal head; increased FDG uptake in previously mentioned heterotopia | FIAS less than 1/month, rare FAS, remote FTBTC | Lamotrigine 200 BID | Right parietal/occipital horn gray matter heterotopia | 12 | 122 | Bilateral MD, bilateral PLV, R ANT |
| 25 | F | 29 | RHD | 21 | Non-lesional; decreased uptake in L lateral temporal lobe (superior temporal gyrus and insula) | FAS/FTBTC once every 2 months | Vimpat 200 mg BID, Keppra mg 1500 BID | L mesial temporal | 17 | 260 | Bilateral ANT, PLV, and MD |
| 26 | F | 50 | RHD | 49 | Hemosiderin staining in right inferior frontal lobe with corresponding subtle decreased FDG uptake | FIAS 1/week, FTBTC 1/year | Vimpat 200 mg BID | R orbitofrontal | 14 | 180 | Bilateral ANT, PLV, and MD |
| 27 | F | 45 | RHD | 3 | Left MTS, no PET available | FAS 1-2/month, rare convulsions | Tegretol ER 600 BID, Zonisamide 200/300, Klonopin 1 mg qHS | Left hippocampus | 13 | 200 | Bilateral MD, only L ANT & L PLV |
